# Regulation of tension-dependent localization of LATS1 and LATS2 to adherens junctions

**DOI:** 10.1101/2025.08.26.672419

**Authors:** Chamika De Silva, Brian A. Kelch, Dannel McCollum

## Abstract

The LIM domain protein LIMD1 is a critical regulator of the Hippo signaling pathway, acting to sequester the kinases LATS1/2 to adherens junctions (AJs) in response to mechanical strain. Here, we identify the molecular basis for LIMD1 binding and recruitment of LATS1/2 to AJs. We show that while the LIM domains of LIMD1 are sufficient for AJ localization and binding to LATS1/2, recruitment of LATS1 to AJ requires both the intrinsically disordered region (IDR) in the N-terminus as well as the LIM domains. We further dissected the LIM domains and found that LIM1 and LIM2, but not LIM3, are necessary for LATS1 AJ localization. Point mutations that disrupt strain sensitivity in either the first or second LIM domain disrupt both binding and recruitment of LATS1/2 to AJs. Mechanistically, LIMD1 binds LATS1/2 through a conserved linear motif, the LATS-LATCH, which we identified by AlphaFold modeling and confirmed by biochemical and localization assays. The LATS-LATCH is both necessary and sufficient for strain-dependent recruitment of LATS1/2 to AJs. Mutation of predicted contact residues within the LATS-LATCH both disrupts its binding to LIMD1 and localization to AJs. These findings define a bipartite mechanism for LIMD1-dependent recruitment of LATS1/2 involving LIM domain-LATCH interactions and N-terminal IDR functions, providing insight into how mechanical signals are transduced through the Hippo pathway.

## INTRODUCTION

The Hippo pathway kinases LATS1/2 regulate the transcriptional co-activator YAP to control cell density dependent inhibition of growth, cell proliferation, apoptosis, stem cell maintenance, and differentiation(1, 2). A major function of LATS1/2 is to control cellular responses to mechanical forces experienced by cells such as changes in cell density, substrate stiffness, shear stress, and tension(3). Pioneering work in *Drosophila*(4) and subsequent studies in mammalian epithelial cells showed that mechanical strain at adherens junctions (AJs) inhibits LATS1/2 by sequestering it at AJs(5, 6). Integral to this regulation is a conserved family of LIM domain proteins related to the *Drosophila* protein JUB that play a central role in strain dependent LATS1/2 regulation(5, 6). LIMD1, the mammalian homolog of the *Drosophila* JUB protein, is essential for strain-dependent regulation of LATS1/2(5, 7). In response to mechanical strain LIMD1 localizes to AJs and recruits LATS1/2. The details of how LIMD1 is recruited to adherens junctions by mechanical strain and stimulated to bind LATS1/2 are poorly understood. Recent work revealed that a subfamily of LIM domain proteins (including LIMD1) recognize strain at AJs(8-10). The LIM domains of these proteins bind F-actin filaments only when it is subjected to mechanical strain, raising the possibility that they could act as tension sensors. In addition, we previously showed that LIMD1 needs to be able to bind strained F-actin in order to associate with LATS1(7). This suggests that binding to strained F-actin somehow stimulates binding to LATS1/2. A key question is how does LIMD1 binding to F-actin under strain trigger binding to and recruitment of LATS1/2 to AJs.

Little is known about how LIMD1 binds to LATS1/2. Therefore, we investigated how LIMD1 interacts with LATS1/2. Through a combination of computational modeling, biochemical, and cell biological experiments we show that the first two LIM domains of LIMD1 bind to a conserved region of LATS1/2 that we call the LATCH. Surprisingly we also find that the non-conserved intrinsically disordered region (IDR) of LIMD1 is required for recruitment of LATS1 to AJs even though it does not directly bind LATS1/2. Together these results suggest possible models for how LIMD1 recruits LATS1/2 to AJs in response to mechanical strain.

## MATERIALS AND METHODS

### Cell culture

Human embryonic kidney cell lines, HEK293, HEK293A and HEK293T were grown in Dulbecco’s Modified Eagle medium (DMEM) (Gibco) supplemented with 10% (v/v) fetal bovine serum (FBS) (Gibco) and 1% (v/v) penicillin/streptomycin (Invitrogen). MCF10A human mammary epithelial cells were cultured in Dulbecco’s Modified Eagle Medium/Nutrient Mixture F-12 (DMEM/F12) (Gibco) supplemented with 5% (v/v) horse serum, 20 ng/mL epidermal growth factor (PeproTech), 0.5 ug/mL hydrocortisone (Sigma), 100 ng/mL cholera toxin (Sigma), 10 ug/mL insulin (Sigma) and 1% (v/v) penicillin/streptomycin (Invitrogen). All cell lines were cultured in a humidified incubator at 37°C with 5% CO_2_. For immunostaining and live cell imaging cells were grown on either 18 mm round cover glasses or µ-Slide 2 well glass bottom chambered coverslips (Ibidi USA), respectively, coated with 0.6 mg/mL collagen (collagen I, rat tail, Gibco) in 20 mM acetic acid solution as previously reported(11).

### Plasmids and molecular cloning

Plasmids for lentiviral production psPAX2 and pMD2.G were a gift from Didier Trono (Addgene plasmid # 12260 and Addgene plasmid # 12259). Lentiviral vector pLJM1-EGFP containing a puromycin resistance gene was used to generate stable cell lines expressing various GFP-fusion proteins. pLJM1-EGFP was a gift from David Sabatini (Addgene plasmid # 19319)(12). Vectors pIC113 and pcDNA3.1(+) (Invitrogen) were used to produce constructs used for transient expression in HEK293 derived cell lines. The pIC113 plasmid was a gift from Iain Cheeseman & Arshad Desai (Addgene plasmid # 44434)(13).

All plasmid constructs used in this study were constructed using the Gibson assembly strategy. Linear DNA fragments encoding genes of interest and vectors with overlapping ends were PCR amplified using Herculase II Fusion DNA Polymerases (Agilent). PCR amplified genes were cloned into appropriate vectors according to the Gibson Assembly Master Mix (New England Biolabs) protocol. Lentiviral plasmid constructs were transformed and maintained in NEB Stable Competent *E. coli* (High Efficiency) (C3040H, New England Biolabs). All other plasmids were transformed and maintained in NEB 5-alpha Competent *E. coli* (High Efficiency) (C2987H, New England Biolabs). All constructs were confirmed by sequencing.

### Lentivirus production and generation of stable cell lines

Lentiviral pLJM1-EGFP constructs, lentiviral packaging plasmid (psPAX2), and the envelop plasmid (PMD2.G) were mixed in a 2:2:1 ratio and transfected into HEK293T cells according to the Effectene Transfection Reagent (QIAGEN) protocol. After 24 hours of transfection, the lentiviral packaging mix was replaced with complete DMEM. The following day the lentivirus containing supernatants were collected and filtered using a 0.45 um pore size filter. MCF10A LIMD1 knockout (LIMD1-KO) cells grown to about 70% confluency were infected with lentiviral supernatant containing 10 ug/mL polybrene in complete DMEM/F12 media for 24 hours. The following day, the media was replaced with complete DMEM/F12 media and incubated until the cell culture reached 90-95% confluency. Transduced cells were selected by passaging in DMEM/F12 media supplemented with 1 ug/mL puromycin (Gibco).

### Cell transfection, immunoprecipitation, and Western blotting

HEK293 or HEK293A cells were transiently transfected for 6 hours using Lipofectamine 2000 Transfection Reagent (Invitrogen) in Opti-MEM (Gibco) media in 6-well tissue culture treated polystyrene plates (CytoOne). Equal amounts of plasmid DNA encoding bait proteins or prey proteins (interacting protein) were transfected into separate wells. Following the 6-hour transfection, media was replaced with complete DMEM media and incubated for 48 hours. Cells were then lysed in immunoprecipitation buffer (10% (v/v) glycerol, 20 mM Tris-HCl, pH=7, 137 mM NaCl, 2 mM EDTA, 1% (v/v) NP-40) supplemented with 1x protease inhibitor cocktail (Sigma-Aldrich), 1 mM phenylmethylsulfonyl fluoride (PMSF), 1 mM sodium orthovanadate and 10 mM sodium fluoride and incubated on ice for 10 minutes. Crude cell lysates were cleared by centrifugation at 15,000 x g at 4°C for 10 minutes and 10% volume from each lysate was saved for input samples. From this point onwards all the steps were performed in a cold room at 4°C. Cleared cell lysates containing bait protein or prey protein were combined in a 2:1 volume ratio and incubated at 4 °C with gentle shaking for 1 hour to allow protein-protein interaction to take place. Combined lysates were then pre-cleared using 50 uL of magnetic Dynabeads Protein G beads (Invitrogen). Immunoprecipitation was carried out using 1 ug of rabbit anti-LIMD1 (Novus biologicals, NBP2-56448) or mouse anti-FLAG antibody (Sigma-Aldrich, F1804) coupled to 50 uL of Dynabeads at 4°C with gentle shaking for 1 hour. Appropriate isotype control antibodies coupled to Dynabeads were used as controls. Immune complexes were separated using a magnetic rack and washed with immunoprecipitation buffer and resuspended in 50 uL of 1 x Laemmli SDS-sample buffer. Both the input and immunoprecipitated fractions were separated by 10% (w/v) SDS-PAGE followed by Western transfer to a nitrocellulose membrane. Membranes were blocked in 5% (w/v) non-fat dry milk dissolved in 0.1% (v/v) Tris-buffered saline with Tween 20 (TBST) and incubated overnight at 4°C with primary antibodies (rabbit anti-LATS2 (1:500, Cell Signaling, 5888S), mouse anti-V5 (1:1000, Cell Signaling, 80076S) or chicken anti-GFP (1:1000, Abcam, ab 13970)).Secondary antibody incubation was done at room temperature for 1 hour using horseradish peroxidase conjugated secondary antibodies (1: 2000). Nitrocellulose membranes were developed using chemiluminescent substrate Clarity Western ECL Blotting Substrate (Bio-Rad) and imaged using Amersham ImageQuant 800 Western blot imaging system. Quantification was done using the ImageJ software and statistical analysis was done using GraphPad Prism 10.5.0 software.

### Fixed-cell Immunofluorescence

MCF10A cells were cultured on glass coverslips coated with 0.6 mg/mL collagen. When cells were about 70% confluent they were fixed with 4% (v/v) paraformaldehyde in UB (UB; 150 mM NaCl, 50 mM Tris pH 7.4) for 10 minutes and permeabilized with 0.5% (v/v) Triton X-100 in UB for 5 minutes at room temperature, then washed 3 times in UB. Cells were then blocked with 10% (w/v) BSA in UB for 1 hour at room temperature and incubated overnight at 4°C with primary antibody (mouse anti-TRIP6 (1:1000, Santa Cruz, sc-365122) and rabbit anti-LATS1 (1:500, Cell Signaling, 3477)). Cells were then washed 3 times and incubated with Alexa Flour-conjugated secondary antibodies for 1 hour at room temperature and washed with UB prior mounting onto glass slides.Coverslips were mounted onto glass slides using Prolong Gold Antifade reagent with 4′,6-diamidino-2-phenylindole (DAPI) (Invitrogen) and left to cure at room temperature overnight. Images were acquired using NIS-Elements-BR software on a Nikon Eclipse Ni-E upright fluorescence microscope using a Plan Apo λ 60x/1.40 oil objective. Images were processed and assembled using ImageJ software.

### Live cell imaging

LIMD1-KO MCF10A cells stably expressing EGFP-tagged proteins were cultured in µ-Slide 2 well glass bottom chambered coverslips (Ibidi USA) coated with 0.6 mg/mL collagen. When cells were about 70% confluent, the GFP signal was observed and images captured using an EVOS M7000 Imaging System (Thermo Fisher Scientific) equipped with an onstage incubator to maintain standard cell culture conditions (37°C with 5% CO_2_ and humidity). Images were acquired with the EVOS M7000 Imaging System software using an Olympus UPlanXApo 40x/0.95 objective. Images were processed and assembled using ImageJ software.

### Drug treatments

MCF10A cells were cultured in collagen coated µ-Slide 2 well glass bottom chambered coverslips. To inhibit tension, cells were treated with DMEM/F12 media supplemented with 25 uM blebbistatin dissolved in dimethyl sulfoxide (DMSO) for 2 hours. As a negative control, cells were treated with DMEM/F12 media supplemented with DMSO. Following drug treatment, media was replaced with fresh DMEM/F12 and cells were imaged using the EVOS M7000 Imaging System.

## RESULTS

### LIMD1-LATS1/2 binding is mediated by the LIMD1 LIM domains but both the LIM domains and IDR region are required to recruit LATS1 to adherens junctions

LIMD1 is required to recruit LATS1 to AJs (5, 7). We investigated how LIMD1 recruits LATS1/2 to AJs in response to mechanical strain. Binding experiments showed that LATS2 bound the carboxy-terminal (aa 468-676) but not the amino-terminal (aa 1-467) region of LIMD1 (**Figure 1A-B**), which is consistent with previous studies showing that LIMD1 related proteins (Ajuba, Zyxin, and TRIP6) bind LATS2 through their 3 tandem LIM domains (6, 14-16) and that the first two LIM domains of LIMD1 are required for it to bind and inhibit LATS1/2(16). To test whether the LIM domains are sufficient to recruit LATS1 to AJs, we used LIMD1 knockout (LIMD1-KO) MCF10A cells(7) to map domains of LIMD1 required for localization and recruitment of LATS1 to AJs. (Note that because endogenous LATS2 is very difficult to detect, all studies involving endogenous proteins focused on LATS1.) For this analysis various LIMD1 deletion mutants were stably expressed in LIMD1-KO cells and assessed for their ability to localize to AJs and to restore localization of endogenous LATS1 to AJs. We first examined the LATS1/2 binding carboxy-terminal LIM domain-containing region (aa 468-676) and the amino-terminal region (aa 1-467) described above. As we showed previously(7), LIMD1-(468-676), but not LIMD1-(1-467), can localize to AJs (**Figure 1D**)(Note that localization of all GFP fusions to LIMD1 (Figure 1-3) (and LATS1/2 (Figure 4-5)) was determined using imaging of live cells since GFP fluorescence did not survive fixation and staining procedures.). Interestingly, both LIMD1-(1-467) and LIMD1-(468-676) are not sufficient for localization of LATS1 to adherens junctions (**Figure 1C**). The carboxy-terminal LIM domains (aa 468-676) are presumably required because they bind LATS1/2 (**Figure 1B**). Why the amino-terminus is required is unclear. The poorly conserved amino-terminus is predicted by AlphaFold to be disordered except for 2 helical regions in the first 68 amino acids. To test whether the helical region in the LIMD1 amino-terminus is important, we deleted the first 94 amino acids, which contains the predicted helical region. This mutant still localized to AJs and was capable of recruiting LATS1 (**Figure 1C**) indicating that it is the intrinsically disordered region of the LIMD1 amino-terminus that is critical for LATS1 recruitment to AJs

**Figure 1.**
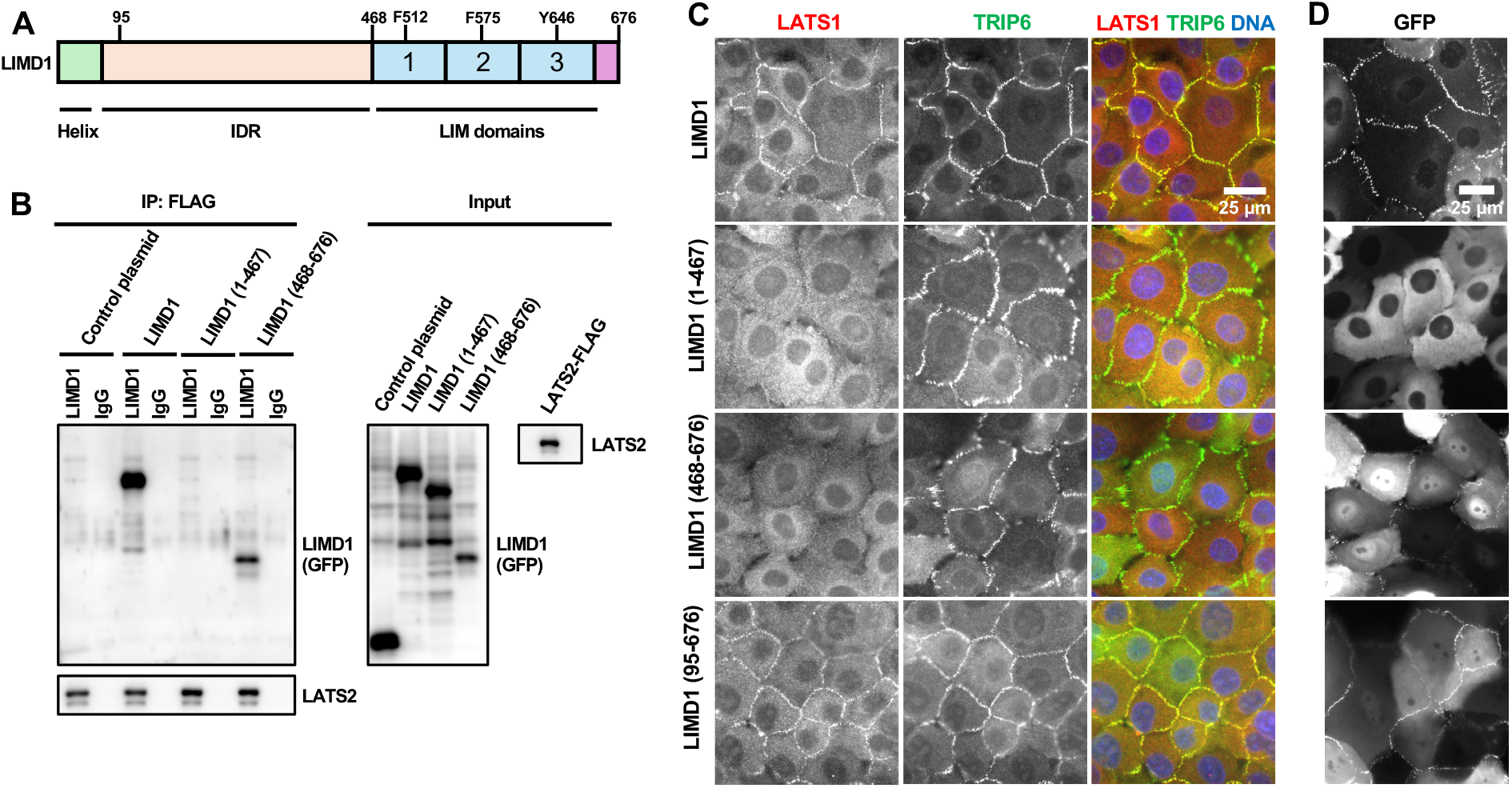
Role of LIM domain and IDR regions of LIMD1 in binding to LATS2 and recruiting LATS1 to adherens junctions. **(A)** Schematic diagram of LIMD1 protein, with amino terminal helical region (Helix), intrinsically disordered region (IDR), LIM domains (1, 2, and 3) indicated. Conserved residues required for binding to strained F-actin (F512, F575 and Y646) are indicated on their respective LIM domains. **(B)** Full-length (LIMD1), the amino-terminal half (1-467), or the carboxy-terminal half (468-676) of LIMD1 were tested for binding to LATS2 by co-immunoprecipitation. GFP (Control plasmid) was used as a control. GFP, GFP-LIMD1 variants, and LATS2-FLAG were separately transfected in HEK293A cells. HEK293A cell lysates transfected with either GFP/GFP-LIMD1 variants or LATS2-FLAG were combined, and anti-FLAG or control (IgG) antibodies were used to isolate immune complexes. Immune complexes and lysates were probed by Western blotting for GFP/GFP-LIMD1 variants and LATS2-FLAG. **(C-D)** LIMD1 knockout (LIMD1-KO) MCF10A cells stably expressing GFP tagged wild-type LIMD1 and LIMD1 variants were established by lentiviral transduction and imaged using fixed and live-cell imaging. **(C)** The indicated cell lines were stained using anti-LATS1 and anti-TRIP6 antibodies. Merged images show LATS1 (red), TRIP6 (green) and DNA (blue). **(D)** The indicated cell lines from part **(C)** were imaged live for GFP fluorescence.

### The first two LIM domains of LIMD1 are required for LATS1 recruitment to adherens junctions

We next examined the requirements for the individual LIM domains of LIMD1 for localization and LATS1 recruitment to AJs. Different combinations of LIM domains were deleted in the context of the full-length protein (**Figure 2A**). Analysis of mutants with just one LIM domain showed that either LIM1 or LIM2, but not LIM3, were sufficient for localization to AJs (**Figure 2C**). However, none of the three single LIM mutants were able to direct LATS1 to adherens junctions (**Figure 2B**). We examined mutants with different combinations of two LIM domains. All combinations (LIM1,2, LIM1,3, LIM2,3) were able to localize to AJs (**Figure 2C**), but only the mutant that had both LIM1 and LIM2 (LIM1,2) was able to recruit LATS1 (**Figure 2B**). Together these results show that both LIM1 and LIM2 are critical for localization to F-actin strain sites and recruitment of LATS1/2 to AJs.

**Figure 2.**
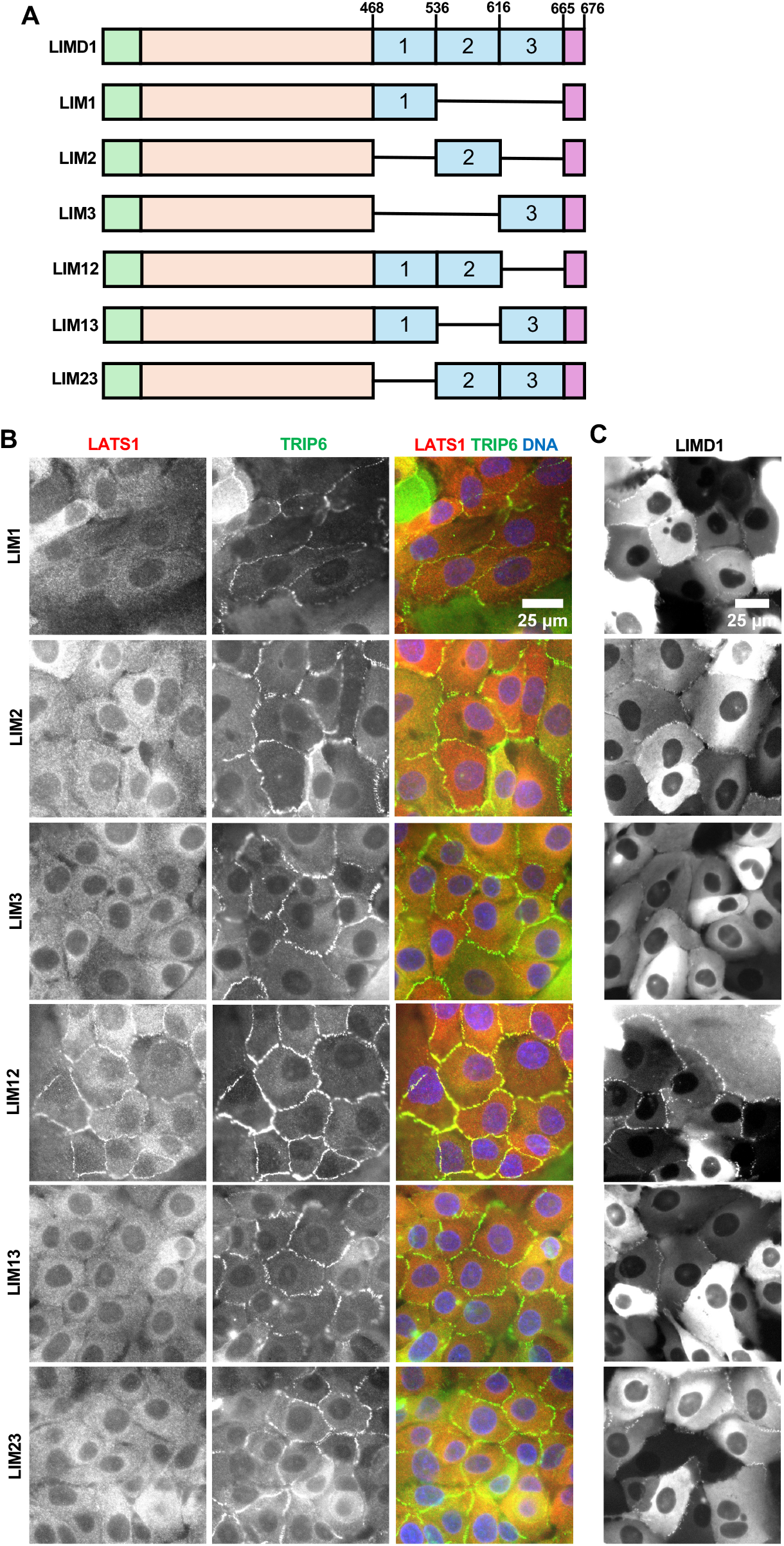
Localization of LIM domain deletion variants of LIMD1 and their ability to recruit LATS1 to adherens junctions. **(A)** Schematic diagram of the different LIM domain deletion variants of LIMD1 used. Each LIM domain is designated by a number (1, 2, or 3). Single horizontal lines denote regions deleted. **(B-C)** LIMD1 knockout MCF10A cells stably expressing GFP tagged LIMD1 deletion mutants shown in **(A)** were established by lentiviral transduction and imaged using fixed and live-cell imaging. **(B)** The indicated cell lines were stained using anti-LATS1 and anti-TRIP6 antibodies. Merged images show LATS1 (red), TRIP6 (green) and DNA (blue). **(D)** The indicated cell lines from part **(C)** were imaged live for GFP fluorescence.

We next examined whether LATS1 recruitment to AJs depends on the ability of the individual LIM domains of LIMD1 to bind to F-actin strain sites. A previous study showed that virtually all LIM domain proteins that bind to F-actin strain sites have a conserved phenylalanine in each LIM domain(8). Mutation of that phenylalanine to alanine in each LIM domain additively disrupts localization to F-actin strain sites(7, 8). The third LIM domain in LIMD1 and several related LIM domain proteins has several distinctive features compared to the other two LIM domains(8). Specifically, it has two additional insertions and lacks the conserved phenylalanine but does have either a phenylalanine or tyrosine (tyrosine in the case of LIMD1) in an adjacent position. We previously showed that mutating all three positions in LIMD1 interfered with its localization to AJs and ability to recruit LATS1 in response to mechanical strain(7). Here we individually mutated the phenylalanine (LIM1 (F512), LIM2 (F575)) or tyrosine (LIM3 (Y646)) to alanine and examined the mutant protein’s ability to localize to AJs and recruit LATS1 (**Figure 3A-B**). Consistent with our LIM domain deletion results, all single mutants were still able to localize to AJs, but mutations in either LIM1 or LIM2 disrupted recruitment of LATS1 to AJs (**Figure 3A-B**). If these mutations specifically affect binding to F-actin strain sites as proposed(8) then our results indicate that both LIM1 and LIM2 must be engaged with strained F-actin to bind LATS1. Alternatively, these mutations may also affect binding to LATS1/2 independent of their effects on binding to F-actin strain sites. Therefore, we assessed how well single LIM domain mutants bound to LATS2. We found that single mutants in LIM1 or LIM2 almost completely disrupted bind to LATS2 (**Figure 3C**). The single mutant in LIM3 impaired LATS2 binding to a lesser extent (**Figure 3C**). That these mutations affect both binding to LATS2 and strained F-actin suggests that they either generally perturb LIM domain structure, or they are directly involved in both binding interactions. Either way they confirm the importance of the first two LIM domains in binding to LATS1/2.

**Figure 3.**
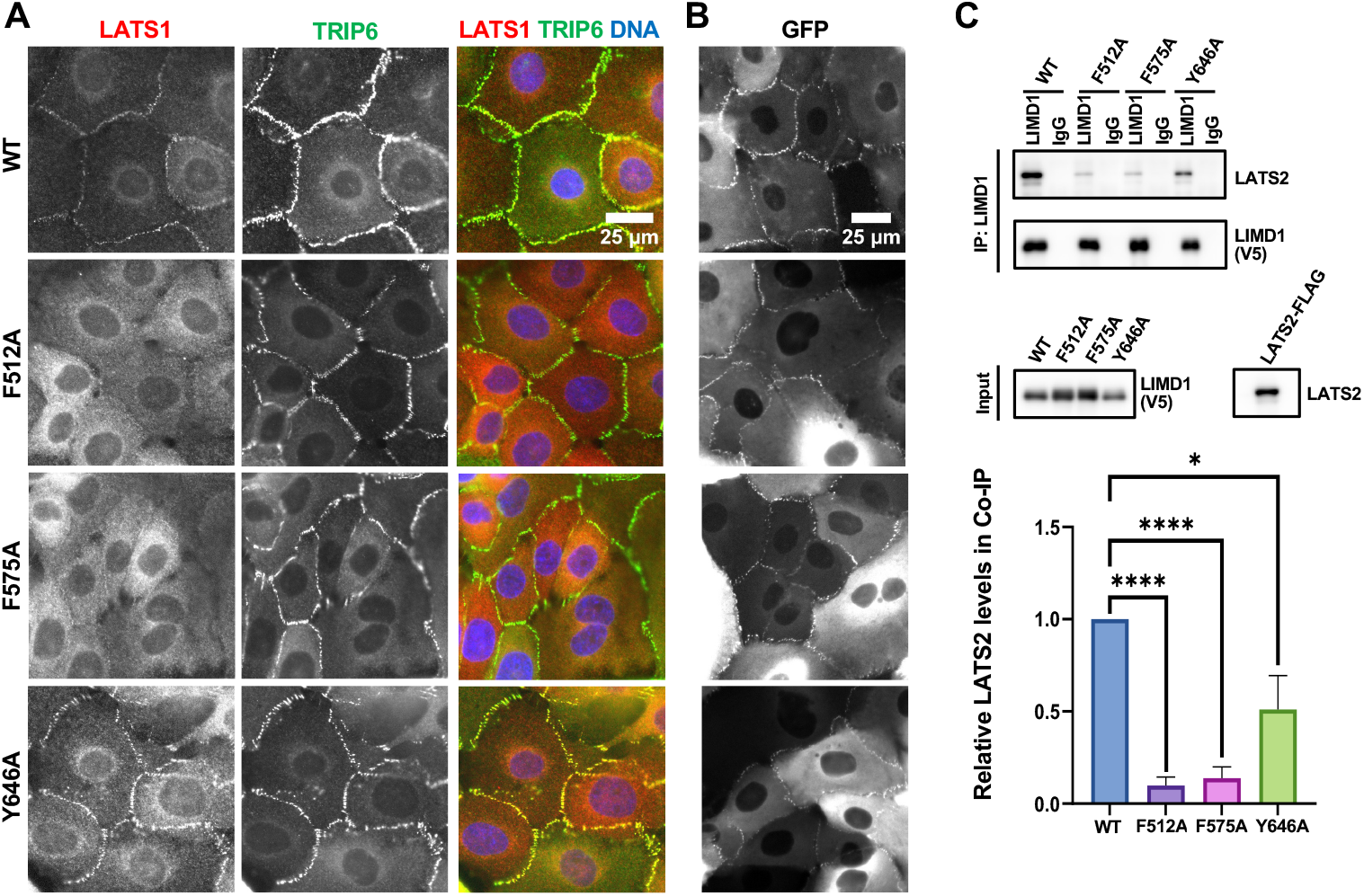
Effect of mechanical strain insensitive LIMD1 mutants on LATS1 recruitment to adherens junctions and LATS2 binding. **(A-B)** LIMD1 knockout MCF10A cells stably expressing GFP tagged LIMD1 or LIMD1 strain insensitive mutants (F512A, F575A, and Y646A, see also Figure 1A) were established by lentiviral transduction and imaged using fixed and live-cell imaging. **(A)** The indicated cell lines were stained using anti-LATS1 and anti-TRIP6 antibodies. Merged images show LATS1 (red), TRIP6 (green) and DNA (blue). **(B)** The indicated cell lines from part **(A)** were imaged live for GFP fluorescence. **(C)** Wild-type (WT) and mechanical strain insensitive mutants (F512A, F575A or Y646A) of LIMD1 were tested for binding to LATS2 by co-immunoprecipitation. V5-tagged LIMD1 mutants and LATS2-FLAG were separately transfected in HEK293A cells. HEK293A cell lysates transfected with either V5-tagged LIMD1 mutants or LATS2-FLAG were combined, and anti-LIMD1 or control (IgG) antibodies were used to isolate immune complexes. Immune complexes and lysates were probed by Western blotting for LIMD1-V5 variants and LATS2. Quantification is shown (mean ± SD; n = 3; *P ≤ 0.05, ****P ≤ 0.0001, t-test).

### LIMD1 binds LATS1/2 through a conserved “LATCH” site

How the LIM domains of LIMD1 bind to LATS1/2 is not known. To address this issue, we first examined which region of LATS1/2 is required for localization to AJs. We found that the largely disordered N-terminus of the LATS1 and LATS2 (aa 1-635 in LATS1, aa 1-598 in LATS2) contained the AJ targeting region (**Figure 4A**). This region overlaps with a region in LATS2 (amino acids 376-660), identified using 2-hybrid screening, as required for binding to the LIMD1 related protein Ajuba(14). To identify potential regions of LATS1/2 that bound to the LIM domains of LIMD1 we used AlphaFold 2/3(17, 18) to model interaction between LIMD1 and this previously identified Ajuba binding region of LATS2 (amino acids 376-660). AlphaFold predicts that the 3 LIM domains of LIMD1 have 2 distinct binding sites in LATS1/2 (**Figure 4B**). The first region (aa431-465 of LATS2) which we term the LATS LATCH is a highly conserve region (**Figure 4C**) that is predicted to run as a linear peptide along the length of the 3 LIM domains, with the highest confidence scores for the regions interacting with LIM domains one and two (**Figure 4B**). The second predicted binding site in LATS1/2 overlaps with the known MOB1 binding site on the LATS1/2(19). This helix turn helix region adjacent to the kinase domain has been termed the helical hairpin(19). This prediction fits with our earlier showing that the LIMD1 related protein TRIP6 can compete with MOB1 for binding to LATS2(6). Of these two regions, the LATS-LATCH is the most important for LIMD1 binding since deletion of the LATCH, but not the helical hairpin, disrupted binding to LIMD1 (**Figure 4D**). In addition, deletion of the LATCH region from the N-terminal regions of both LATS1 and LATS2 disrupted their localization to AJ (**Figure 4A**).Furthermore, the GFP-LATS2-LATCH alone was sufficient for binding to LIMD1 (**Figure 5A**). The LATS2-LATCH region alone fused to GFP localized to AJ in a manner that depended on mechanical tension (**Figure 5B-C**). A LATS2-LATCH mutant targeting four amino acid residues (4mut: L441G, V444A, K445D, R448D) predicted by the AlphaFold2 structure as important contact residues with LIM2 of LIMD1 fails to localize to adherens junctions (**Figure 5B**) and is unable to bind LIMD1 (**Figure 5A**). Together these results support the AlphaFold model showing that the LIMD1 LIM domains interact with the LATCH region of LATS2 (and likely LATS1).

**Figure 4.**
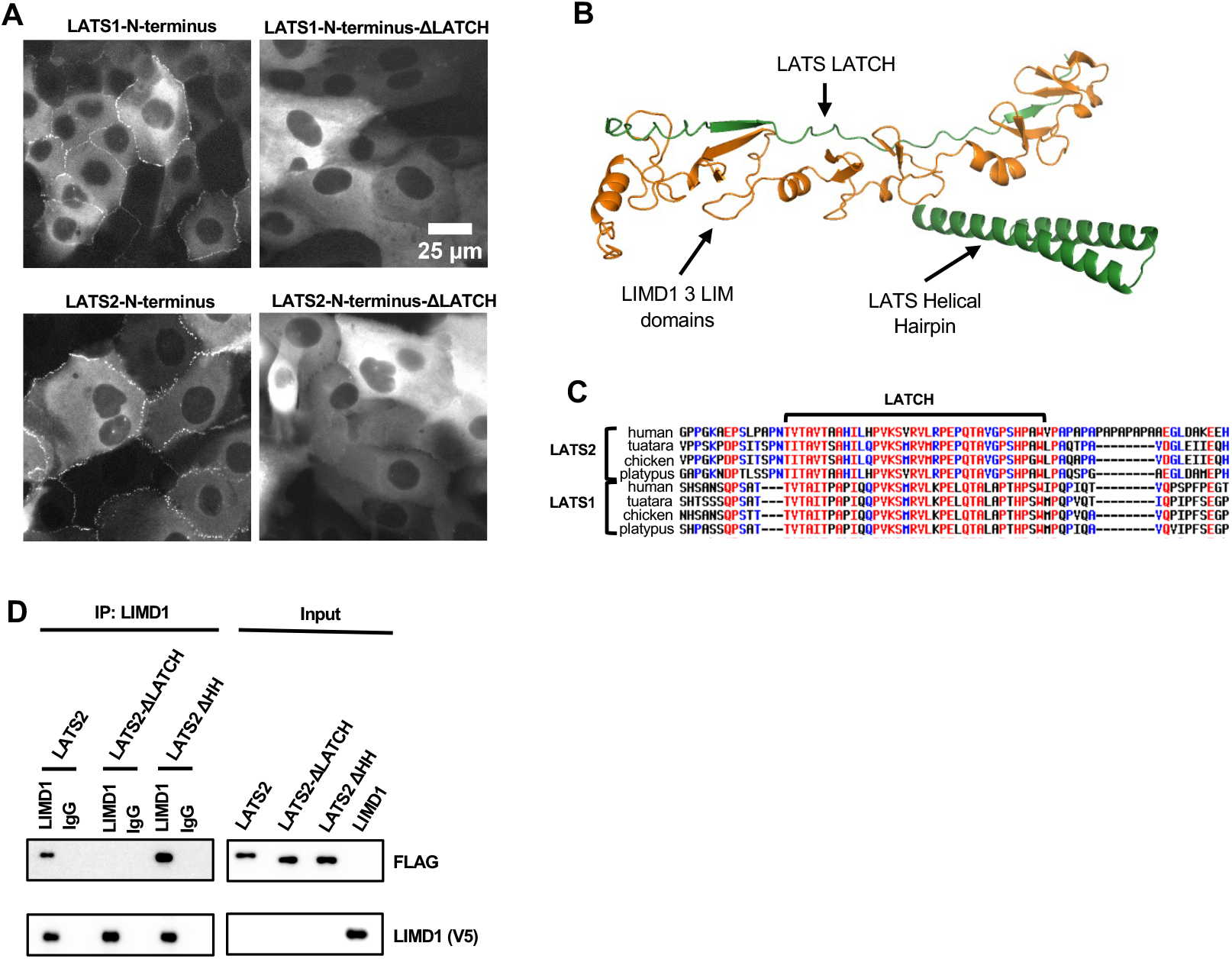
LATS-LATCH is necessary for LATS1/2 recruitment to AJs and for LATS2 binding to LIMD1. **(A)** Live-cell imaging of MCF10A cells stably expressing GFP tagged LATS1 or LATS2 N-terminal regions (aa 1-635 in LATS1, aa 1-598 in LATS2) with and without (ΔLATCH) the LATCH regions (aa 472-520 in LATS1, aa 419-467 in LATS2) as indicated. **(B)** AlphaFold2 model showing the three tandem LIM domains of LIMD1 (green) and two regions of LATS2 (orange) that are predicted to interact (LATS-LATCH and the Helical Hairpin). **(C)** Multiple sequence alignments of LATS1/2 from the indicated species showing the conserved LATCH region. **(D)** Full-length LATS2 (LATS2) or LATS2 with either the LATCH deleted (LATS2-ΔLATCH, aa 419-467 deleted) or with the helical hairpin region deleted (LATS2-ΔHH, aa 599-667 deleted) were tested for binding to LIMD1 by co-immunoprecipitation. FLAG-LATS2 variants and LIMD1-V5 were separately transfected in HEK293 cells. HEK293A cell lysates transfected with either V5-tagged LIMD1 or LATS2-FLAG mutants were combined, and anti-LIMD1 or control (IgG) antibodies were used to isolate immune complexes. Immune complexes and lysates were probed by Western blotting for LIMD1-V5 variants and FLAG-LATS2.

**Figure 5.**
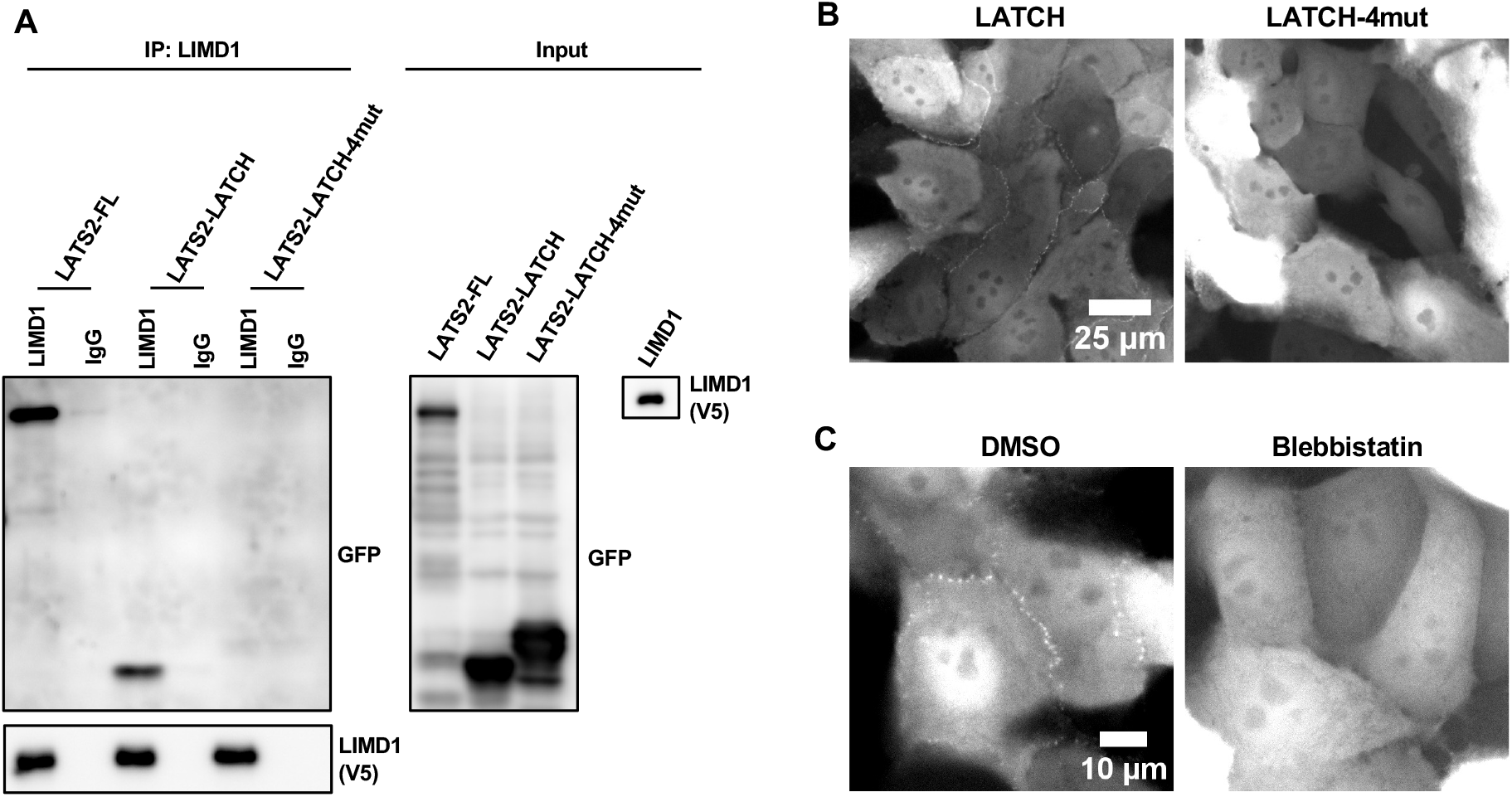
The LATS-LATCH is sufficient to bind to LIMD1 and localize to AJs in a tension-dependent manner. **(A)** Full-length LATS2, LATS2-LATCH and LATS2-LATCH-4mut, were tested for binding to LIMD1 by co-immunoprecipitation. Lysates were prepared from HEK293A cells separately transfected with GFP-tagged LATS2, LATS2-LATCH, LATS2-LATCH-4mut, or LIMD1-V5. Lysates from cells expressing LIMD1-V5 were mixed separately with those expressing the different GFP-LATS2/LATCH constructs, and anti-LIMD1 or control (IgG) antibodies were used to isolate immune complexes. Immune complexes and lysates were probed by Western blotting for GFP (LATS2/LATCH) and V5 (LIMD1). **(B)** Live-cell imaging of MCF10A cells stably expressing GFP tagged LATS2-LATCH or LATS2-LATCH-4mut. (C) Live-cell imaging of MCF10A cells stably expressing GFP tagged LATS2-LATCH treated with (Blebbistatin) or without (DMSO) Blebbistatin.

## DISCUSSION

It has been known for many years that the Hippo pathway kinases LATS1/2 are regulated by mechanical strain(3). Although strain-dependent regulators of LATS1/2, such as LIMD1, have been identified, the molecular mechanisms governing how strain is sensed and transduced into regulation of LATS1/2 have remained elusive. To gain insight into this question, we investigated the molecular basis for the mechanical strain-dependent interaction between LIMD1 and LATS1/2. As a first step towards understanding why LIMD1-LATS1/2 association at AJs is strain sensitive we defined the region of each protein required for this interaction. AlphaFold modeling provided critical testable hypotheses for how the proteins interacted by predicting that a linear peptide from LATS1/2 (the LATCH) binds along the length of the three LIM domains of LIMD1, with the highest confidence scores for interaction of the LATS-LATCH with the first two LIM domains of LIMD1. The AlphaFold model also suggested a potential interaction between the LIM domains and the helical hairpin region of LATS1/2. The helical hairpin has been shown to bind the LATS1/2 activator MOB1 to promote LATS1/2 activating autophosphorylation(19). Our experiments have confirmed several aspects of this model, in particular the interaction between the first two LIMD1 LIM domains and the LATS-LATCH. We observe that the first two LIM domains of LIMD1 are required for binding and recruitment of LATS1/2 AJs. Second, the LATCH sequence is necessary and sufficient for binding to LIMD1 and recruitment of LATS1/2 to adherens junctions. Third, mutations in predicted interface residues in the LATCH disrupt binding to LIMD1 and localization to AJs. The other potential binding site for LIMD1 on LATS1/2 (the helical hairpin) was not required for LIMD1-LATS1/2 binding. This is consistent with the predicted LIMD1-LATCH interface having ∼4.5 fold greater sized interface than the LIMD1-helical hairpin interface. However, there is some reason to suspect that the predicted LIMD1-helical hairpin interaction may be relevant in vivo. For example, we previously showed that TRIP6, a LIMD1 related protein, competes for binding to LATS2 with MOB1, a LATS1/2 activator that binds the helical hairpin. One possibility could be that the LATCH acts as a strain-dependent tether to allow the weaker LIMD1-helical hairpin interaction to compete for binding with MOB1. Further biochemical experiments will be needed to test this possibility.

A key question is why binding of the LATCH to LIMD1 depends on mechanical strain in vivo. Although we can detect binding between LIMD1 and LATS1/2 in vitro, we suspect that the in vitro binding is weak since we can only detect it with overexpressed proteins. We think that binding of the LIM domains to strained F-actin filaments somehow makes them better at binding the LATS-LATCH sequence. Possible models include the LIM domains undergoing a conformational change when bound strained F-actin that enhances binding to the LATS1/2 LATCH or alternatively binding to strained F-actin may bring the individual LIM domains into register to enhance binding to the LATCH sequence. Another model could be that when LIMD1 binds strained F-actin the LATCH actually binds both the LIM domains and the strained F-actin filament. In all these models a high affinity binding site for the LATCH would only be created when LIMD1 is bound to strained F-actin. Our observation that strain-insensitive LIMD1 mutations in LIM1 (F512) and LIM2 (F575) impair LATS1 recruitment to adherens junctions, suggests that binding to strained F-actin is required for this process. However, because these mutants also have defects in binding LATS2 in vitro, interpretation of the mutant phenotype is complicated. The LATS2 binding defect of the F512A and F575A mutants suggests that these residues could contribute to binding to both strained F-actin and LATS1/2. Alternatively, these mutations could have more general effects on the folding of the LIM domains. Together these results indicate that some caution is warranted when assuming that these mutations solely affect binding to strained F-actin.

One of the surprising findings of this work is that the IDR region of LIMD1 is required for recruitment of LATS1 to adherens junctions. This result is surprising because the LIM domains of LIMD1 are sufficient both for binding to LATS1/2 and localization of LIMD1 to AJs. How the LIMD1 IDR region promotes LATS1 localization to AJs is not known. One intriguing possibility is that the IDR region of LIMD1 may promote liquid-liquid phase separations (LLPS) that are important for LATS1 accumulation at AJs. This would be consistent with previous work showing that the IDR region of LIMD1 is important for LLPS formation and recruitment of proteins to focal adhesions(20). LLPS has been implicated in other aspects of Hippo pathway signaling(21-27). Further studies will be needed to test whether LIMD1 driven LLPS formation drives LATS1 localization to AJs.

In summary, this study uncovers the molecular mechanism by which the mechanosensitive protein LIMD1 recruits the Hippo pathway kinases LATS1/2 to AJs in response to mechanical strain. Using domain mapping, mutagenesis, and AlphaFold modeling, our work shows that LIMD1 uses its first two LIM domains to bind LATS1/2 via a conserved “LATS-LATCH” motif in the LATS N-terminus. These LIM domains also interact with strained F-actin, suggesting that mechanical tension enhances LATCH binding, potentially through conformational changes or a three-way binding interface between LIMD1, LATS1/2, and strained F-actin. Our surprising discovery that LIMD1’s IDR is also required for LATS1 recruitment raises the possibility that the IDR promotes liquid–liquid phase separation to concentrate signaling components. Together, the findings show how LIMD1 recruitment of LATS1/2 to AJs integrates direct protein-protein binding with cytoskeletal engagement to provide new insight into how mechanical forces regulate Hippo signaling at cell-cell junctions.

## ACKNOWLEDGEMENTS

This research was supported by National Institutes of Health (grant RO1 GM058406).

